# A Framework for Accurate Prediction of Plastic-Degrading Enzymes using Convolutional Neural Networks

**DOI:** 10.1101/2024.10.20.619257

**Authors:** Soharth Hasnat, Fariah Anjum Shifa, Shabab Murshed, Sarker Tanveer Ahmed Rumee, MST Murshida Mahbub

**Affiliations:** Depertment of Genetic Engineering and Biotechnology, East West University, Dhaka, Bangladesh; Department of Computer Science and Engineering, University of Dhaka, Bangladesh

**Author notes:** **Correspondence to:** Sarker Tanveer Ahmed Rumee, and MST Murshida Mahbub. Authors equally contributed to this work (Soharth Hasnat, Fariah Anjum Shifa).

**Keywords:** Plastic-degradation, machine-learning, pollution, enzymes and sequence

## Abstract

The growing accumulation of plastic waste presents a significant environmental challenge, necessitating innovative approaches to mitigate its impact. Enzymatic degradation has emerged as a promising solution for addressing plastic pollution. However, the isolation and characterization of plastic-degrading enzymes (PDEs) through laboratory experiments are costly, time-consuming, and often complicated by nonculturable microorganisms. Consequently, accurate in silico identification of PDEs is desirable to explore the diversity of natural enzymes and harness their potential for combating plastic pollution. This study introduces a novel feature extraction strategy for identifying plastic-degrading enzymes, incorporating Autocorrelation (AAutoCor), Composition of k-spaced Amino Acid Pairs (KSAP), Dipeptide Deviation from Expected Mean (DDE), Composition/Transition/Distribution (C/T/D), Conjoint Triad, and Secondary Structure. A combination of ANOVA and XGBoost, feature selection methods, was applied to optimize the feature dimensions for improved performance. Seven supervised machine learning models were employed to evaluate the dataset: Convolutional Neural Network, Random Forest Classifier, Feedforward Neural Network, Logistic Regression, Naive Bayes Classifier, K-nearest Neighbor, and XGBoost Classifier. Among these models, the CNN model demonstrated the best performance, achieving an accuracy of 0.96, an F1 score of 0.80, and an ROC-AUC score of 0.96. These findings underscore the potential of the proposed system as an accurate predictor of plastic-degrading enzymes from environmental sequences. This approach significantly enhances efforts to develop sustainable solutions to plastic waste by accelerating the discovery of novel PDEs.

## 1. Introduction

Plastic waste continuously poses threats to the soil, marine, and freshwater environments.^1,2^ When broken down, plastics may convert into micro- and nano-plastics, which may enter the human body through ingestion or inhalation with significant health risks. Despite their harmful effects, the use of plastic is increasing daily along with growing populations.^3^ Annually, over 0.3 billion tons of plastics are produced worldwide, and only 21% have been recycled; the rest is released to the environment.4 To date, over 200 enzymes have been reported with the plastic degradation capacity.^5,6^ The already discovered plastic degrading enzymes offer promise in identifying more such enzymes through computational approaches. Mining the huge sequence databases that may harbour undiscovered degrader sequences is timely and compelling as they may include superior features. Although laboratory-based experiments are continuously finding plastic degrading enzymes, they are time consuming and costly.^7–9^ While wet lab-based findings are the solid verification of the biological activities, computational screening can save time and costs which made the later approaches as routine procedures to predict a particular function before conducting lab-based experiments. Among the computational approaches, machine learning-based approaches are gaining popularity in finding protein functions from the sequence data. This study applied several machine learning methods using wet lab-verified plastic degrading enzymes as positive control compared with proven non-plastic degrading enzymes as negative control to get a highly accurate plastic degrader predictor. The organization of the paper is as follows. Section 2 describes the background information necessary for the study. In section 3 we discuss the methods and parameters of model used for the implementation. The experiment results and future works are presented in section 4.

## 2. Background

### 2.1. Performance metrics

#### 2.1.1. F1-score

F1-score is utilized to quantify binary classification systems, which classify examples into ‘positive’ or ‘negative.’ It is the harmonic mean of the model’s precision and recall.^10^ The metric is calculated using the following formula:

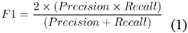

#### 2.1.2. Matthews Correlation coefficient

The Matthews Correlation Coefficient (MCC), also called the phi coefficient, serves as an evaluative metric for binary classification models. It offers a balanced evaluation of a model’s effectiveness by considering true positives, true negatives, false positives, and false negatives. The MCC is determined by utilizing the following formula:

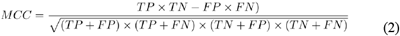

Here: TP, TN, FP, and FN refer to True Positives, True Negatives, False Positives, and False Negatives respectively. MCC is very useful for dealing with imbalanced datasets or evaluating the overall quality of a binary classification model. So, it is considered in our evaluation for balancing the trade-off between precision and recall.

#### 2.1.3. ROC curve

A Receiver Operator Characteristic (ROC) curve is a graphical plot that shows the diagnostic ability of binary classifiers.^11^ It thoroughly analyzes the model’s behavior when a range of threshold values are considered, depicting the compromise between specificity and sensitivity. The Area Under the ROC Curve (AUC) encapsulates the interplay between these rates, concisely summarizing the model’s overall performance. We utilized this metric to inform our decision-making process regarding the model’s acceptability.

## 3. Method and materials

### 3.1. Training Dataset

#### 3.1.1. Positive dataset

Two hundred and eight PD enzyme sequences were collected from the Plastic Degradation Database (Plastic DB) to create a positive dataset.^5^ We further cleaned up the dataset by discarding 26 proteins that contained non-standard amino acids. Finally, the positive dataset contained 182 protein sequences (S1 File). The gathered PD proteins varied in length from 108 to 914.

#### 3.1.2. Negative dataset

1523 non-PD protein sequences were extracted from UniProt to assemble the negative dataset.^12^ We constructed a search model using an enhanced search option to find proteins that are not involved in the degradation activities. To do that, the names of twenty known PD enzymes were removed from the search model. Later, manual verification was performed to ensure that proteins only involved in the synthesis processes were included in the negative dataset. The supplementary file (S2 File) contains the constructed search model.

### 3.2. Feature extraction

Six descriptors have been chosen to extract features from the PD and non-PD enzymes (Table 1), and they are used to extract features from biological sequences.^13–17^ This process used Biopython and ftrCOOL package (an RStudio package). ^13,18^

**Table 1.** Descriptors of feature extraction methods.

| Descriptors | No. of features | Packages | Extracted features |  |
| --- | --- | --- | --- | --- |
|  |  |  | Positive Dataset | Negative Dataset |
| Autocorrelation (AAutoCor) | 240 | ftRCOOL | 43,680 | 365,520 |
| Composition of K-spaced amino Acids pairs (kSAP) | 3600 | ftRCOOL | 655,200 | 5,482,800 |
| Composition/Transition/Distribution (C/T/D) | 343 | ftRCOOL | 28,126 | 522,389 |
| Conjoint triad | 343 | ftRCOOL | 28,126 | 522,389 |
| Dipeptide deviation from the expected mean (DDE) | 400 | ftRCOOL | 32800 | 609,200 |
| Helix, turn, and sheet | 03 | Biopython | 246 | 4,569 |
| Total |  |  | 788,178 | 7,506,867 |

#### 3.2.1. Autocorrelation

Autocorrelation is used to extract features based on the distribution of amino acid properties in the protein sequence.^19^ To get the feature, we added three autocorrelation descriptors, Geary, Moran, and Normalized Moreau-Broto (NMBroto), in the R script. Subsequently, eight Aaidx (Amino acid index) IDs were incorporated into the script. Finally, the script was run with package ftrCOOL to acquire the feature matrix. The script is available in the supplementary file (S3 File).

#### 3.2.2. Composition of k-Spaced Amino Acids pairs (KSAP)

This descriptor calculates the frequency of all amino acid pairs with k spaces.^13^ In the R script, we added the range (rng) vector as a number, where each vector element shows the number of spaces between amino acid pairs. For each k in the rng vector, a new vector (whose size is 400) was created containing the frequency of pairs with k gaps. In our analysis, we set the rng value to ten (S3 File), which provided 3600 features for each protein in the feature matrix.

#### 3.2.3. Conjoint Triad (CT)

We employed this descriptor to explore each amino acid’s triad properties in protein sequences. For the calculation, this function turns 20 amino acids into seven classes according to their dipoles and volumes of the side chains. CT descriptor counts any three continuous amino acids as a unit resulting in 343 features extracted from each sequence.^13,20^ This function returns a feature matrix corresponding to the R script (S3 file).

#### 3.2.4. Composition/Transition/Distribution (C/T/D)

C/T/D is a group of descriptors representing the amino acid distribution pattern based on the protein’s precise structural and physicochemical properties. Seven types of physical properties have been taken to calculate these features: hydrophobicity, normalized Van der Waals volume, polarity, polarizability, charge, secondary structures, and solvent accessibility.^13^ This group of descriptors can calculate each sequence’s composition, transition, and distribution, and we use these parameters as vectors. Each element of the vector shows the number of spaces between the first and the second amino acids and the second and third amino acids of the tripeptide.^13^ For each k in the rng vector, a new vector (size: 73) is created, which contains the frequency of tri-amino acid with k gaps (S3 file).

#### 2.3.5. Dipeptide Deviation from Expected mean (DDE)

DDE is considered one of the most critical descriptors for developing a sequencebased predictor for accurately identifying proteins.^17,21^ Dipeptide composition has been widely adopted in various protein function prediction methods, such as Bcell linear epitope prediction.^22,23^ For biological research purposes, the dipeptide combination of a protein sequence is primarily calculated as:

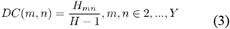

Where Hmn is the quantity of paired mn amino acids, and H expresses the size of the protein sequence. Next, the theoretical mean (TM) and theoretical variance (TV) of a protein sequence are computed as:

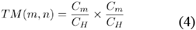

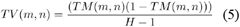

where Cm is the number of codons, encrypting for the first amino acid, and Cs is the number of codons, encrypting for the second amino acid in the given dipeptide. Finally, DDE is intended based on TV, TM, and DC as:

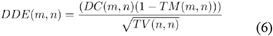

#### 3.2.6. Feature of secondary structure (helix, turn, and sheet)

Helix, turn, and sheet are the principal structural features of proteins.^24^ The secondary structures of plastic-degrading enzymes are expected to possess unique features. Hence, we extract quantitative features from the protein sequences by targeting the three crucial features (helix, turn, and sheet) using Biopython.^18^

### 3.3. Feature elimination methods

#### 3.3.1. Analysis of variance (ANOVA)

Our analysis determined the statistically significant difference between the means of three or more independent groups using ANOVA. It selects features by comparing the average of the dependent variable across different categories of each independent variable. This indicates that the independent variable significantly influences the dependent variable when there are substantial differences in the averages.1 Additionally, ANOVA aids in ranking the identified features according to the computed F-statistic.^25,26^

#### 3.3.2. ANOVA and Permutation Feature Importance

An ensemble of feature selection strategies was employed, integrating ANOVA with Permutation Feature Importance (PFI). This approach involves the union of features selected by ANOVA and those identified through PFI, capitalizing on the strengths of both methodologies. ANOVA provides insight into the statistical significance of features, while PFI ^27^ evaluates the impact of each feature on model performance by measuring the degradation in performance when feature values are permuted. This combination of methods ensures that features are statistically significant and critical to maintaining high model performance, resulting in a more robust and reliable feature set.

#### 3.3.3. ANOVA and XGBoost Feature Selection

Another feature selection method employed was the integration of ANOVA with eXtreme Gradient Boosting (XGBoost).^28^ XGBoost, a gradient boosting technique, assigns scores to features based on their contribution to improving model predictions across multiple decision trees.^29,30^ Combining ANOVA’s statistical significance with XGBoost’s predictive capability ensures that the features selected are statistically significant and highly influential in enhancing model accuracy. This hybrid approach provides a comprehensive and optimized set of features, thereby improving the overall performance and reliability of the predictive model.

### 3.4. Feature set/ descriptor set creation

Creating diverse feature combinations in machine learning can significantly enhance both model performance and interpretability. Each feature captures distinct aspects of the data, and their fusion can influence key performance metrics such as accuracy, precision, and recall. Through experiments comparing individual features and their combinations (Table 2), we observed that combining features uncovers deeper patterns in the data, leading to improved performance. To identify the optimal feature set, we constructed and evaluated five distinct feature sets using various machine-learning algorithms. Feature Set 1 includes Composition, Transition, and Distribution (CTD), Autocorrelation (Corr), Dipeptide Deviation from Expected Mean (DDE), K-spaced amino acid pair (KSAP), and Secondary Structure (SS). Feature Set 2 comprises CTD, Conjoint Triad (CT), DDE, KSAP, and SS. Feature Set 3 consists of CTD, CT, DDE, Corr, and SS. Feature Set 4 contains KSAP, CT, DDE, Corr, and SS. Finally, Feature Set 5 includes KSAP, Conjoint Triad (CJT), CTD, Corr, and SS. By evaluating the performance of these feature sets, we aim to determine which combination best captures the underlying data patterns and leads to the highest model accuracy.

**Table 2.** Comparison of individual features and feature set performance.

| Feature Name | Acc | F1 | PR AUC | ROC AUC | MCC |
| --- | --- | --- | --- | --- | --- |
| AutoCorrelation | 0.94 | 0.71 | 0.78 | 0.78 | 0.7 |
| Secondary Structure | 0.90 | 0.58 | 0.63 | 0.73 | 0.54 |
| C/T/D | 0.94 | 0.72 | 0.89 | 0.78 | 0.72 |
| DDE | 0.95 | 0.78 | 0.82 | 0.83 | 0.76 |
| KSAP | 0.94 | 0.74 | 0.80 | 0.81 | 0.73 |
| CJTriad | 0.94 | 0.72 | 0.75 | 0.81 | 0.69 |
| Feature Set | 0.95 | 0.78 | 0.84 | 0.90 | 0.75 |

### 3.5. Application of machine learning algorithms

A total of seven different models, namely Convolutional Neural Network, Random Forest Classifier, Naive Bayes Classifier, XGBoost Classifier, K-Nearest Neighbor, Logistic Regression, and Basic Neural Network, were incorporated into this study. Each model technique was applied to the filtered dataset, but the Convolutional Neural Network (CNN) performed better considering the performance.

The CNN model was selected for the classification task in our investigation due to its superior accuracy compared to other machine-learning techniques. It used the feature sets as the input sequence feature vector. Five convolutional layers, three fully connected hidden layers, and a fully connected output layer with two output neurons that contain sigmoid activations for binary classifications make up the architecture of our deep neural network. Each convolutional layer has a kernel size of 1, a leakyReLU activation function, and batch normalization. The 1D filter performs correlation operations on the matrix. Since Sigmoid and Tanh can lead to the vanishing gradient problem, ReLu was utilized as the activation function. Batch normalization stabilizes the training by normalizing the internal covariate shift in each layer and reducing it.^31^

Additionally, it introduces some noise to lessen the overfitting of the data. A variety of filters are included in the convolution layer. Zero-padding is used in all convolutional layers to produce an output length equal to the input length. Unless otherwise stated, max-pooling with an identical window size and stride follows each convolutional layer. The fully connected hidden layer uses one hundred twenty-eight units with ReLU activation functions. Dropout, through random neuron deactivation during training, encourages the exploration of diverse paths and feature combinations. This enhances the model’s capacity to generalize to unseen data, a critical aspect in developing an accurate system.^32^

In the training phase, a dropout probability of 0.2 was employed for the hidden layers as this value seemed to perform the best among other values. Regularizers are techniques used to prevent overfitting and make models more interpretable.^33^ Common kernel regularizers such as L1 and L2 regularization add penalty terms to the loss function to promote smaller weights in the model. After experimenting with various values, we used the L1 kernel regularizer with a strength of 0.001. All the parameters were initialized using He initialization,^34^ and we trained the model for 74 epochs.

After each epoch, the parameters were updated using TensorFlow’s implementation of Adam optimizer with a loss function of binary cross entropy. Three fully connected layers are then flattened, followed by a max-pooling layer. In CNN architectures, max pooling is frequently employed to minimize the spatial dimensions of feature maps while keeping crucial information. We used a sigmoid activation function in the output layer to determine the likelihood of each class label for classifying plastic-degrading enzymes.

### 3.6. Synthetic minority oversampling technique (SMOTE)

Our datasets have 182 samples of positive class and 1492 samples of negative class, which shows the necessity of using an oversampling technique to balance the data. Classification in an imbalanced dataset is challenging because of highly skewed data, with one class (the minority class) having significantly fewer examples than the other class (the majority class). This scenario can pose a bias towards the majority class, leading to a suboptimal result. We employed the SMOTE (Synthetic Minority Oversampling Technique) technique for synthesizing new samples to balance the data, thereby creating a more balanced distribution and improving the model’s ability to learn from the minority class. We used the sampling strategy ‘minority’ to create new instances of minority class and balance the class distribution.35

## 4. Result and discussion

### 4.1. Feature Selection

When we observed the six individual features with enormous dimensions, it became evident that dimension reduction was necessary to filter out the most impactful ones. Retaining unnecessary features can increase dataset noise, adversely affecting model performance. It is important to note that there is no universal feature selection method; the choice of technique depends on the specific characteristics of the data. Consequently, three feature elimination methods were evaluated, as described in Section 3.3.

Before selecting an appropriate feature selection method for the dataset, we created five feature sets by combining the six individual features, as detailed in Section 3.4. Through rigorous analysis of the performance of these feature sets across the three distinct feature selection methods(table 3, it was observed that the ANOVA+XGBoost method consistently outperformed the other two methods across all five feature sets. A previous machine learning approach on plastic degrading enzymes only applied the XGB technique to select essential features.^36^

**Table 3.** Performance Comparison of Different Feature Selection Methods.

| Feature Set | Method | Accuracy | F1 | ROC | Matthews | Pr auc |
| --- | --- | --- | --- | --- | --- | --- |
|  | ANOVA | 0.962 | 0.78 | 0.83 | 0.72 | 0.82 |
| Feature set 1 | ANOVA + Permutation | 0.960 | 0.79 | 0.84 | 0.731 | 0.83 |
|  | ANOVA + XGB_feature_selection | **0.961** | **0.795** | **0.85** | **0.735** | **0.84** |
|  | ANOVA | 0.95 | 0.76 | 0.81 | 0.75 | 0.81 |
| Feature set 2 | ANOVA + Permutation | 0.960 | 0.78 | 0.84 | 0.77 | 0.82 |
|  | ANOVA + XGB_feature_selection | **0.96** | **0.8** | **0.85** | **0.77** | **0.83** |
|  | ANOVA | 0.96 | 0.79 | 0.83 | 0.731 | 0.83 |
| Feature set 3 | ANOVA + Permutation | 0.961 | 0.8 | 0.84 | 0.734 | 0.84 |
|  | ANOVA + XGB_feature_selection | **0.96** | **0.8** | **0.85** | **0.74** | **0.84** |
|  | ANOVA | 0.95 | 0.78 | 0.84 | 0.75 | 0.82 |
| Feature set 4 | ANOVA + Permutation | 0.96 | 0.79 | 0.84 | 0.74 | 0.82 |
|  | ANOVA + XGB_feature_selection | **0.96** | **0.79** | **0.87** | **0.76** | **0.84** |
|  | ANOVA | 0.96 | 0.77 | 0.82 | 0.73 | 0.83 |
| Feature set 5 | ANOVA + Permutation | 0.96 | 0.79 | 0.85 | 0.74 | 0.82 |
|  | ANOVA + XGB_feature_selection | **0.96** | **0.79** | **0.85** | **0.77** | **0.85** |

The integration of XGBoost^37^ with ANOVA effectively leveraged the strengths of both techniques, resulting in more discriminative feature selection compared to the other two methods, thereby improving model performance. Moreover, the application of the ANOVA+XGBoost feature selection method led to significant dimension reduction in the features Autocorrelation, DDE, CTD, KSAP, SS, and Conjoint Triad, as shown in Table 4. The dimension of the Autocorrelation feature was reduced from 240 to 136, DDE from 400 to 108, CTD from 148 to 38, KSAP from 3600 to 109, and Conjoint Triad from 343 to 137. This dimension reduction was crucial in mitigating potential overfitting in the models and simplifying computational processes. The decrease in overfitting risk, facilitated by dimension reduction, ensured the model’s accuracy.

**Table 4.**
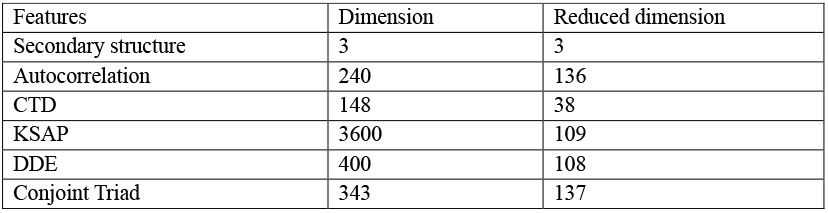
Dimension reduction of features.

### 4.2. Oversampling

Previous studies stated that imbalanced datasets could lead to biased models since the algorithms favor the class with more instances.^38^ In this work, imbalanced data can lead to suboptimal model performance by favoring the majority class while struggling to identify the minority class. Our oversampling techniques increased the number of instances in the minority class to match the majority class, ensuring balance among the different classes.^39^ There are various methods to perform oversampling. After careful evaluation, the SMOTE (Synthetic Minority Oversampling Technique) method was considered best for this investigation.^35^

A comparative study was conducted between models trained on oversampled data and those trained without sampling. The aim was to investigate the influence of oversampling on the model’s generalization ability and its effectiveness in predicting both classes. The analysis output (supplementary file 5) demonstrated the superiority of oversampled datasets. Models trained on oversampled data consistently

### 4.3. Model performance analysis

In this study, we evaluated the performance of seven models-Convolutional Neural Network (CNN), Random Forest Classifier, Naive Bayes Classifier, K-Nearest Neighbor, Logistic Regression, XGBoost Classifier, and a Basic Neural Network—on five feature sets, designated as F1, F2, F3, F4, and F5, to determine which model produces the best results.

Through a detailed analysis of performance metrics across all model and feature combinations, we concluded that feature set 4, which includes Secondary Structure(SS), Conjoint Triad(CT/CJT), DDE, Autocorrelation (AutoCorr), and K-spaced Amino Acid Pair (KSAP), performed as the most critical set of features for classifying plastic-degrading enzymes (Table 5). The Convolutional Neural Network (CNN) model trained on feature set 4 demonstrated superior performance compared to other model-feature combinations. Figure 3 shows the performance of CNN models trained on five feature sets consecutively.

**Table 5.**
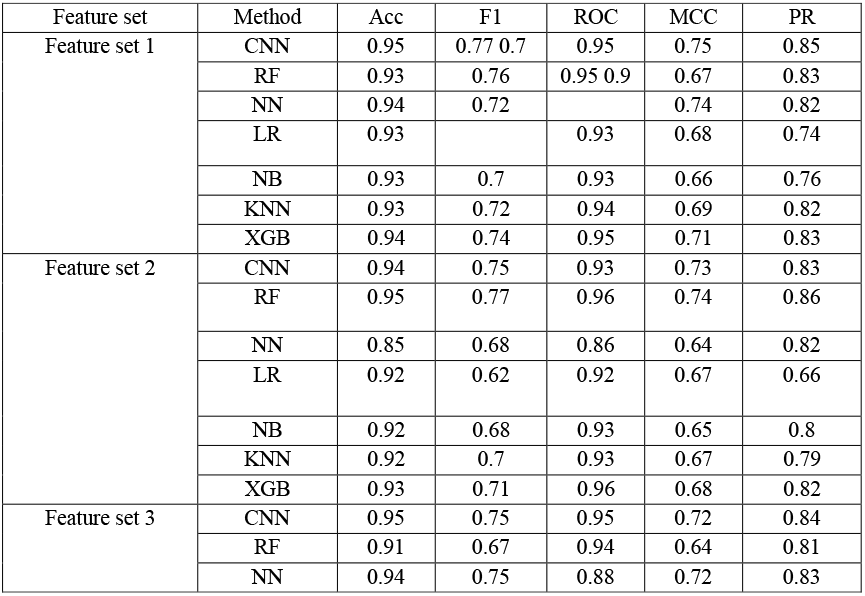
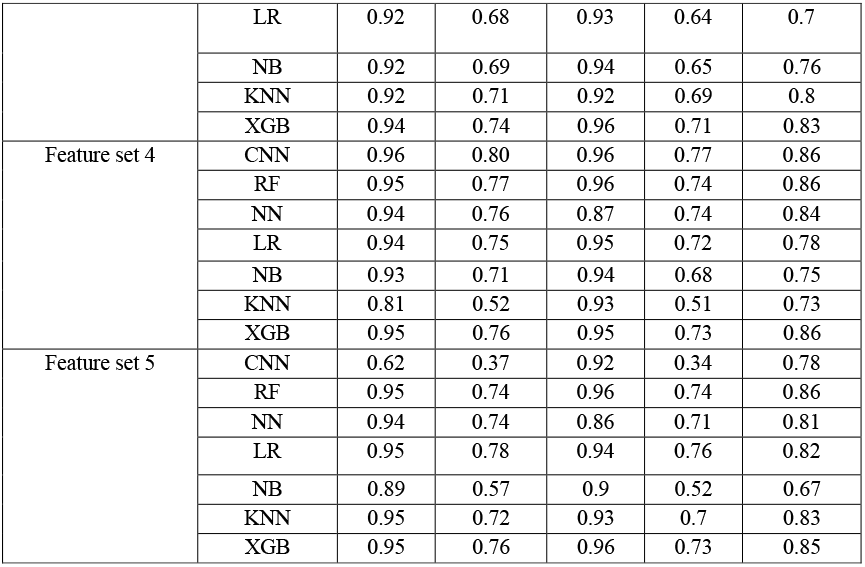
Performance analysis of five feature set combinations using five methods.

**Fig. 1.**
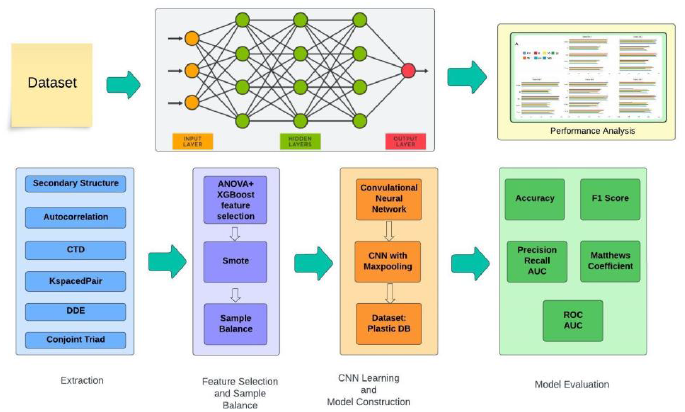
Graphical abstract represents the workflow that is utilized in this research.

**Fig. 2.**
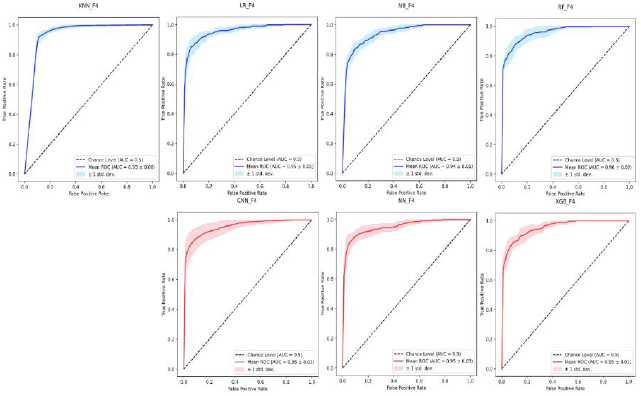
Performance analysis of ROC curves for each model on feature set 4

**Fig. 3.**
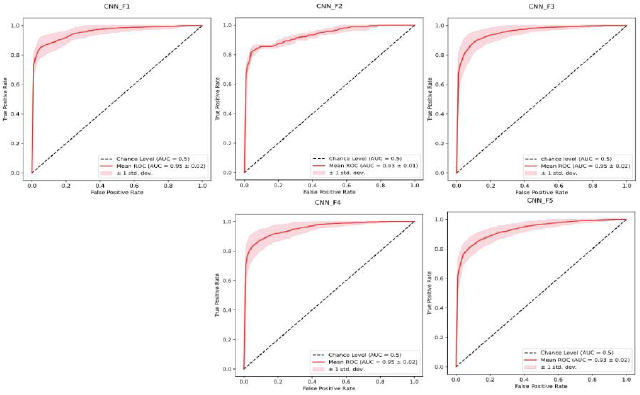
CNN model performance comparison on five feature sets

**Fig. 4.**
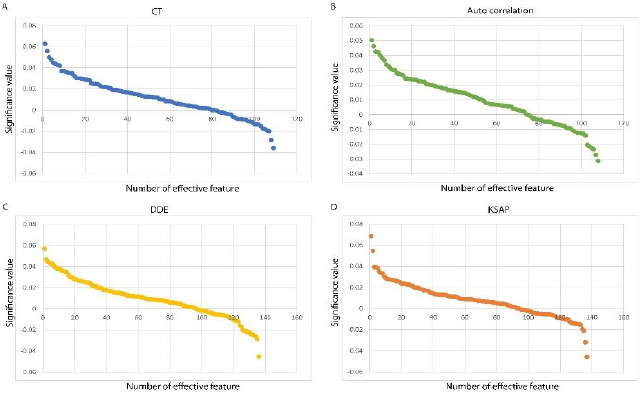
Identification of essential features involved in plastic degradation. A. Importance of top feature types of CT on the model performance B. Importance of the 136 features of Auto correlations and their impact on model performance C. Assessment of the 108 features of DDE and their influence on model performance D. Impact of the 108 features of K-spaced amino acid pairs on model performance.

While accuracy only considers correctly categorized tuples unaffected by their classes (plastic-degradable/non-plastic-degradable), we place greater weight on F1 due to its balance of precision and recall. The results, as shown in Table 5, indicate that CNN trained on feature set 4 consistently outperforms the other variations of the model and feature sets. While some models achieved F1 scores ranging from 57% to 77%, CNN surpassed them with a score of 80%. In conclusion, our investigation demonstrated that CNN outperforms XGBoost and other models in classifying plastic-degrading enzymes, showcasing the power of deep learning techniques in handling complex biological data.

The primary objective of this research was to accurately identify plasticdegrading enzymes (PDEs) from protein sequences using an in-silico approach. A key finding of our study is identifying critical features that significantly contribute to plastic degradation. By employing a novel combination of ANOVA and XGBoost (XGB) feature selection, we pinpointed four main feature categories that impact model performance: K-spaced amino acid pairs (KSP), Dipeptide Deviation from Expected (DDE), Autocorrelation, and Composition, Transition, Distribution (CTD). Through our analysis, Autocorrelation emerged as the most influential feature set. Model performance dropped significantly when Autocorrelation was excluded from the training process, as indicated in Supplementary File S4. Following this, we found KSP, CTD, and DDE features necessary. To further assess feature importance, we categorized them into two groups based on their contribution to the F1 score: positive values, which caused a decrease in the F1 score when removed, and negative values, where feature removal resulted in a slight increase in the F1 score. The graph in Supplementary File S4 illustrates these positive and negative values. While features with positive values were deemed crucial for distinguishing plastic-degrading enzymes, those with negative values—despite slightly enhancing the score—were minimal in effect and thus excluded from further analysis. Future work will focus on several key areas to improve model performance further. These include expanding the dataset with more diverse protein sequences, exploring deep learning models such as Transformers for better feature extraction, and addressing class imbalance with more advanced oversampling techniques. Additionally, incorporating domain knowledge on protein structure and function may enhance the biological interpretability of the model’s predictions.

## 5. Conclusion

In conclusion, our research presents a novel approach for accurately identifying plastic-degrading enzymes through machine learning. By carefully selecting and evaluating features that play a critical role in distinguishing these enzymes, we have laid the groundwork for more efficient enzyme discovery in the fight against plastic pollution. Future improvements in model architecture and dataset expansion will further enhance the potential of this approach.

## Supporting information

https://doi.org/10.6084/m9.figshare.27263676

## Author’s Contributions

**Soharth Hasnat:** Formal analysis, Conceptualization, Methodology, Software, Visualization, Writing – review & editing. **Fariah Anjum Shifa:** Formal analysis, Methodology, Software, Visualization, Writing – review & editing. **Shabab Murshed:** Methodology, Software, Visualization, Writing – review & editing. **Sarker Tanveer Ahmed Rumee:** Conceptualization, Supervision, Validation, Writing – review & editing. **M Murshida Mahbub:** Conceptualization, Supervision, Writing – review & editing.

## Author’s approval

**All authors reads and gives consent for publication**

## Supplementary Information

All supplementary are available at https://doi.org/10.6084/m9.figshare.27263676

